# Larger Pupils are Associated with Improved Visual, but not Auditory, Near-Threshold Detection

**DOI:** 10.64898/2026.07.29.741499

**Authors:** Veera Ruuskanen, Sebastiaan Mathôt

**Author notes:** Address for correspondence: Veera Ruuskanen, Department of Experimental Psychology, University of Groningen, Grote Kruisstraat 2/1, 9712TS Groningen, The Netherlands.

## Abstract

Prestimulus pupil size is associated with near-threshold detection performance in both the visual and auditory domain, a relationship that is commonly attributed to arousal. However, given that larger pupils also let more light into the eye, in the visual domain this relationship is likely also driven by optical effects. To better understand this, we investigated how pupil size, skin conductance, and electroencephalographic (EEG) measures relate to detection performance in both a visual and an auditory task. We found that larger pupils were associated with higher sensitivity in the visual condition but lower sensitivity in the auditory condition. Skin conductance was negatively related to visual sensitivity, but unrelated to auditory sensitivity. EEG power measures were not related to sensitivity in either condition, though pupil size was positively correlated with alpha and beta power. Together, these results suggest that in visual detection the relationship between pupil size and performance is driven by both arousal and optics, whereas in auditory detection the relationship is driven solely by arousal. More broadly, our findings highlight the pupil as an active and functional component of the visual system.

---

Sensory perception is influenced by moment-to-moment fluctuation in the state of the brain and sensory organs. This is especially evident at the perceptual threshold, where the same stimulus is sometimes perceived and sometimes not. Previous research has found that spontaneous fluctuations in pupil size and in electroencephalographic (EEG) power are associated with changes in near-threshold detection performance both in the visual and auditory domain (Beerendonk et al., 2024; Koenig & He, 2025; Pilipenko et al., 2026; Podvalny et al., 2021; Ruuskanen et al., 2025). These effects are typically attributed to fluctuations in arousal (reflected in pupil size) and cortical excitability (reflected in EEG). However, these accounts overlook another potential cause: the optical consequences of pupil-size changes. The size of the pupil regulates the amount of light that enters the eye, thus maintaining a balance between visual sensitivity and visual acuity (Mathôt, 2020; Ruuskanen & Mathôt, 2026a; Vilotijević & Mathôt, 2024). In the visual domain this likely partly drives the relationship between pupil size and detection performance (Ruuskanen et al., 2025; Ruuskanen & Mathôt, 2026a). Here, we address this possibility by examining the effects of pupil size and other arousal-related measures on detection performance in the visual and auditory domain.

In visual detection tasks, medium-to-large pre-stimulus pupil size is consistently associated with improved performance (Beerendonk et al., 2024; Eberhardt et al., 2022; Ruuskanen et al., 2025; Ruuskanen & Mathôt, 2026b). The effect is robust to variation in low-level stimulus parameters such as color and eccentricity (Ruuskanen & Mathôt, 2026b), also holds for higher-level stimuli like objects and faces (Koenig & He, 2025), and is present in a dual-task setup (Claeys et al., 2026). In our view, the relationship between pupil size and visual detection performance is driven by two underlying mechanisms. One mechanism is related to optics: larger pupils allow more light to enter the eye, thus improving visual sensitivity, at the expense of acuity (Ruuskanen & Mathôt, 2026a; Vilotijević & Mathôt, 2024). This optical effect drives a linear (or at least monotonically increasing) relationship between pupil size and detection performance.

Another mechanism is related to fluctuations in arousal, driven by the activity of subcortical nuclei including the locus coeruleus (LC), the dorsal raphe, and the superior colliculus (SC) (Grujic et al., 2024; Joshi et al., 2016). These structures are causally involved in both pupil dilation and arousal-regulation via the release of neurotransmitters such as norepinephrine (NE) and acetylcholine (ACh) (Grujic et al., 2024; Reimer et al., 2016). The effect of arousal drives an inverted-U shaped relationship between pupil size and performance. In other words, performance is generally best at intermediate levels of arousal and deteriorates at low and high levels, as characterised by the classical Yerkes-Dodson curve (Yerkes & Dodson, 1908). This canonical relationship is observed in detection tasks using visual, auditory, and tactile stimuli (Beerendonk et al., 2024; de Gee et al., 2024; McGinley et al., 2015; Podvalny et al., 2021; Schriver et al., 2018). However, there is some context-dependent variation in the shape of the relationship. Specifically, in the visual domain results are more mixed, with the relationship between pupil size and performance sometimes manifesting as an inverted-U (Beerendonk et al., 2024), other times as rather linear (Eberhardt et al., 2022; Mathôt & Ivanov, 2019), and yet other times as a combination of the two (Ruuskanen et al., 2025). Presumably, this variability reflects the relative contribution of arousal, which drives an inverted-U relationship, and optics, which drives a linear relationship, to task performance (Ruuskanen & Mathôt, 2026a).

Since pupil size is strongly modulated by arousal, teasing apart the optical effect and arousal effect on visual detection performance is non-trivial. One way to do so is to introduce another measure of arousal that is also associated with detection performance, such as power in the alpha (8-12Hz) and beta (13-30 Hz) bands of the EEG. Alpha power is generally associated with cortical excitability (a lowered threshold for neural firing), such that lower alpha power signals higher excitability (Lange et al., 2013; Romei et al., 2008) and seems to be similarly inversely correlated with arousal (Barry et al., 2020; Schubring & Schupp, 2021). When it comes to visual detection, alpha power is often inversely related to hit rate, such that lower pre-stimulus power is associated with increased hit rate (Ergenoglu et al., 2004; Koenig & He, 2025). However, the relationship between alpha and visual detection performance is not consistent across studies, with some findings suggesting that there is no relationship (Boncompte et al., 2016; Ruuskanen et al., 2025) or that alpha *phase* (but not power) may play a larger role (Busch et al., 2009; Pilipenko et al., 2026). Beta power has been studied less, especially in the context of visual detection. However, the patterns seem to be similar, with beta power being inversely correlated with arousal (Schubring & Schupp, 2021) and detection performance (Koenig & He, 2025). Overall, while the relationship between EEG power and detection performance is not straightforward, there is some evidence pointing towards alpha and beta suppression being associated with changes in detection rates.

Despite both measures being related to arousal and to detection performance, the effects of pupil size and EEG power have largely been studied in isolation. The picture that emerges from the few recent studies that have been conducted is somewhat unclear. We have found that the effect of pupil size on detection performance is not mediated by EEG power in either the alpha, theta, or beta band (Ruuskanen et al., 2025), while others have found that alpha power does mediate the effect of pupil size (Koenig & He, 2025). Some studies indicate that pupil size affects detection sensitivity (d’) while alpha power affects the decision threshold (the criterion) (Pilipenko & Samaha, 2024). However, effects of pupil size on response tendency and criterion have also been shown (Nuiten et al., 2026; Podvalny et al., 2021; Ruuskanen et al., 2025; Ruuskanen & Mathôt, 2026b). Furthermore, many studies have found pupil size to correlate positively with hit-rate while alpha and beta power both correlate negatively (Koenig & He, 2025; Podvalny et al., 2021) In sum, there is no clear understanding of the exact interplay of pupil size and EEG power in determining detection performance, but there is some evidence pointing towards the effects being dissociable.

Even so, it remains unclear whether EEG power suppression and pupil dilation reflect the same underlying arousal mechanisms. If they do not, the observed dissociation is not in itself evidence for distinct optical and arousal mechanisms—arousal itself could consist of (at least) two dissociable components, one expressed in EEG power and one expressed in pupil dilation. This ambiguity stems from how arousal is defined and operationalised. As discussed above, pupil size as a marker of arousal is based on subcortical neuromodulatory activity, most notably in the LC-NE system (Joshi et al., 2016). However, it has been suggested that arousal is a multi-dimensional concept, encompassing not only cognitive arousal (which is generally measured with pupil size), but also emotional, sexual, and general physiological arousal, among others (Sabat et al., 2025). On the other hand, the inverse relationship between EEG power and arousal has been established in studies that manipulated arousal via emotionally charged or sexual imagery (Schubring & Schupp, 2021). Notably, pupil size and EEG power in both the alpha and beta bands are often positively correlated both during task performance and at rest (Mathôt et al., 2023; Montefusco-Siegmund et al., 2022; Podvalny et al., 2021; Ruuskanen et al., 2025; Waschke et al., 2019). Given that larger pupils are thought to reflect increased arousal, this seems to contradict the assumed inverse EEG power-arousal relationship. This discrepancy may be a reflection of the multidimensional nature of arousal. While there is thus some overlap between EEG power, pupil size, and arousal, the exact nature of the relationship remains unclear. This is important for the current study, because it means that previous studies examining both measures in visual near-threshold detection have not fully addressed the question of whether and how the optical effects of pupil size affect detection performance. Here we extend previous research by simultaneously examining the visual and auditory domain.

Pupil size has also been linked to performance differences in the auditory domain, where the observed relationship generally closely resembles an inverted-U shape (Beerendonk et al., 2024; Doll et al., 2025; McGinley et al., 2015; Murphy et al., 2011). This means that detection is best at intermediate pupil sizes, reflecting intermediate arousal. This relationship seems to be less variable than what is observed in visual detection (although at least one study also reports a linear relationship; Doll et al., 2024). Presumably, this is because in auditory tasks there is no performance boost associated with the optical consequences of increased pupil size, and the relationship between pupil size and performance is fully governed by arousal.

Here, we aim to further investigate the optical effects of spontaneous pupil-size fluctuations in visual detection. During a near-threshold detection task consisting of a visual and an auditory condition, we will concurrently measure pupil size, EEG, and skin-conductance. Skin-conductance is another physiological measure that has been linked to arousal variation and pupil size (Chang et al., 2025; Wang et al., 2018). We expect to find a positive correlation between pupil size and skin-conductance in both conditions. To our knowledge no previous studies have examined the relationship between skin-conductance and detection performance, and thus, we do not formulate explicit hypotheses in this regard. However, given its relation to arousal, skin-conductance should theoretically exhibit an inverted-U relationship with performance.

Regarding EEG, we expect to find a positive correlation between pupil size and power in the alpha and beta bands, and a negative correlation between broadband EEG power and visual detection accuracy, replicating previous findings. Given the paucity of studies investigating the relationship between EEG power and auditory detection performance, we do not formulate specific hypotheses. To extend previous research on the relationship between arousal-related EEG markers and near-threshold detection, we also include an analysis of 1/f slope. This aperiodic component in the EEG is thought to reflect the excitation and inhibition balance and has been suggested to function as another marker of arousal (Lendner et al., 2020).

Finally, we expect to find different relationships between pupil size and performance in the two conditions, such that in the visual condition the relationship is relatively linear, while in the auditory condition it more closely resembles an inverted-U.

## Methods

### Open-practices statement

Experimental materials, raw data, and analysis scripts are available on the open-science framework (https://osf.io/8dw3b). The study was not pre-registered.

### Statement on the use of AI

In the process of this research, generative AI (https://sigmundai.eu/) was used to assist in writing parts of the analysis scripts. All AI generated code was reviewed by the authors before use and no content created solely by AI is included in the final manuscript.

### Participants

A total of 30 participants with normal or corrected-to-normal vision participated in the experiment. In the absence of predicted effect sizes, we did not conduct a power analysis, but based our sample size on our laboratory standard of 30 participants. In the case of corrected-to-normal vision, participants wore contact lenses (not glasses) during recording. All participants were undergraduate students at the University of Groningen and were compensated for their participation with partial course-credit. After receiving information about the study in both verbal and written form, participants provided written informed consent prior to participation. Five participants were discarded because the staircase in the auditory condition failed to maintain accuracy at around 75%, causing the auditory task to be either too difficult or too easy (accuracy < 55% or > 90%). One participant was discarded during processing due to excessive eye movements, leading to more than 50% unusable data. Data processing steps and trial exclusion criteria are detailed below. A total of 24 participants were included in the final analysis. The study was approved by the Ethics Committee of the Psychology Department at the University of Groningen (study code: PSY-2324-S-0420).

### Detection task

Participants completed a detection task consisting of a visual and an auditory condition, with the order of conditions counterbalanced between participants. The structure of blocks and trials was identical across conditions. The conditions only differed with regard to the detection targets, which are described below (in the sections *visual stimuli* and *auditory stimuli*). The experiment and stimuli were created with OpenSesame (version 4.0, *Melodramatic Milgram*) (Mathôt et al., 2012) using the PsychoPy (Peirce et al., 2019) backend with PyGaze (Dalmaijer et al., 2014) for eye tracking. The experiment was presented on a 27” LCD monitor with a refresh rate of 120 Hz and a resolution of 1920 × 1080 pixels, at a viewing distance of approximately 60 cm.

Each trial began with the presentation of a circular gray (1.4 cd/m^2^; RGB = 48, 48, 48) fixation dot, which was maintained on the screen throughout the trial. The size of the dot was 0.43 degrees of visual angle (dva) (15 px). The background of the screen was dark grey (7.7 cd/m^2^; RGB = 96, 96, 96).

Total trial length varied between 4000 and 5000 ms, randomly determined. Targets were present on 50% of the trials, with the exact moment of presentation varied to avoid temporal preparation effects. Targets could not be presented in the first 1000 ms or the last 50 ms of a trial. At the end of the trial the fixation dot transformed into a question mark to indicate that a response should be given. Participants were required to press the right arrow key if they had detected a target and the left arrow key if they had not. The task did not progress until a response was given. Responses were given with the index and middle fingers of the right hand. The trial progression is depicted in Figure 1.

**Figure 1.**
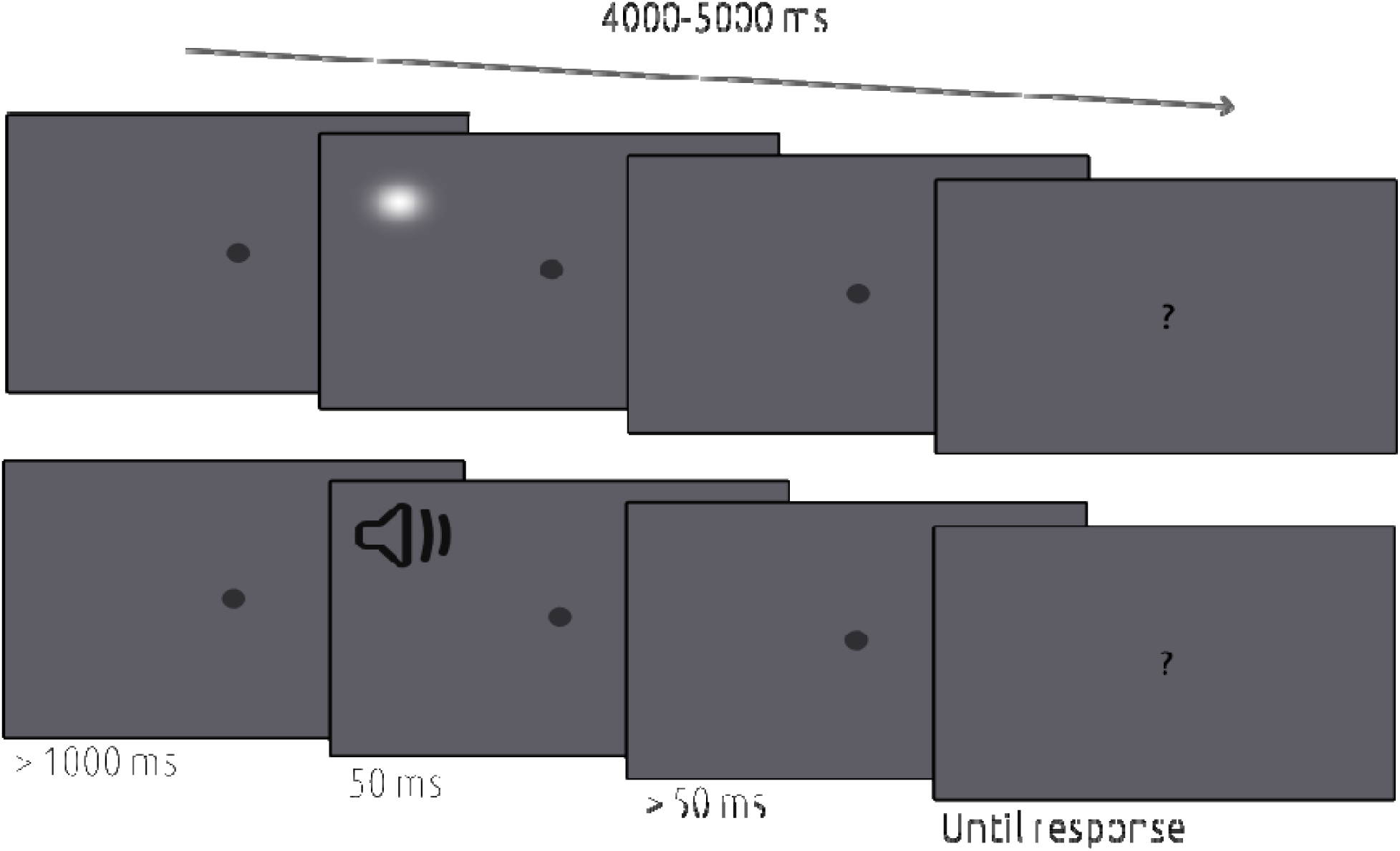
The detection task. The upper row depicts the visual condition and the lower row depicts the auditory condition. Total trial length varied between 4000 and 5000 ms and targets could not be presented in the first 1000 ms or the last 50 ms. Targets were present on 50% of the trials in both conditions. Responses were required for the task to continue.

Each condition consisted of 12 blocks. The first, fifth, and ninth blocks were staircase blocks, consisting of 50 trials. During these blocks the opacity (visual condition) and volume (auditory condition) of the targets was adjusted with a QUEST adaptive-staircase procedure (Watson & Pelli, 1983) to maintain approximately 75% accuracy (details of staircase procedures can be found below in *visual stimuli* and *auditory stimuli*). Trials in staircase blocks were not included in the analysis. The remaining nine blocks consisted of 40 trials each. Thus, there were a total of 510 trials in each condition, with 360 included in the analysis. Furthermore, each condition began with 10 practice trials, during which the staircase was also running. Blocks were separated with self-paced breaks, and the two conditions were separated by a longer break (minimum duration three minutes), with participants being encouraged to take more time if needed.

#### Visual stimuli

The detection targets in the visual condition were luminance patches, defined as instances of a PsychoPy (Peirce et al., 2019) GratingStim with a Gaussian mask and one color (white; max. 84.8 cd/m^2^; RGB = 255, 255, 255).

As mentioned above, target opacity was determined with a QUEST adaptive-staircase procedure (Watson & Pelli, 1983). The starting value of the staircase was set to 0.1 and the standard deviation of the threshold estimate was 0.25.

Targets were presented in the periphery of the visual field. Target position was determined at the beginning of each trial by drawing a random angle between 0 and 360, and transforming it into x and y coordinates on a circle with radius of 14.49 dva (500 px). Targets were flashed for 50ms.

#### Auditory stimuli

The detection targets in the auditory condition were beeps, defined as instances of a PsychoPy (Peirce et al., 2019) Synth stimulus, The frequency of the target sounds were 1000Hz and they were played for 50ms.

As mentioned above, target volume was determined with a QUEST adaptive-staircase procedure (Watson & Pelli, 1983). The starting value of the staircase was set to 0.1 and the standard deviation of the threshold estimate was 0.25.

### EEG acquisition

The EEG signal was recorded from 26 Ag/AgCI scalp electrodes positioned according to the international 10-20 system, as well as six external electrodes, placed on the mastoids and above and below the eyes. The signal was recorded at 1000 Hz using a TMSI REFA 32 amplifier controlled by OpenViBE data acquisition software (Renard et al., 2010). During acquisition the signal was referenced to the average of all electrodes and offline it was re-referenced to the mastoids. EEG was measured continuously.

### Pupil size measurement

Pupil size was measured with an Eyelink 1000 eye tracker with a 35mm lens (SR Research Ltd., 2022) monocularly from the left eye only. The tracker was calibrated at the beginning of the experiment with a 9-point calibration procedure. Pupil size was measured from the beginning to the end of each trial. Participants rested their head on a chinrest, keeping the eye-to-monitor distance fixed at 60cm. The chinrest also limited head movements that could cause muscle artifacts in the EEG signal or cause the eyetracker to lose the eye.

### Skin conductance measurement

Skin conductance was measured with two connected Ag/AgCI electrodes placed on the index and middle fingers of the left hand. The electrodes were connected to the same TMSI REFA 32 amplifier as the EEG cap and other external electrodes and the signal was also recorded at 1000 Hz using the OpenViBE acquisition software (Renard et al., 2010).

### Procedure

Participants were given information about the study in both verbal and written form upon arrival at the lab. After receiving all the necessary information, they signed an informed consent form. After the informed consent procedure, the EEG cap was placed on the participants head, such that electrode Cz was positioned in the middle of the nasion and inion. The cap was then filled with gel and optimised until all electrode offsets were low (< 20 kΩ) and stable. Next, the skin conductance electrodes were taped on the fingers and the eyetracker was calibrated.

After preparation participants completed the task, as described above. Task instructions were given verbally during the informed consent procedure and again on the screen at the beginning of the task. Participants were tested in a Faraday cage with constant, dim (<1 Lux) illumination. The whole experimental session, including preparation, task performance and debriefing, took approximately 2.5 hours per participant.

### Data processing

All data processing was done with custom scripts written in Python, using the package eeg-eyetracking-parser (Mathôt et al. 2023), which relies on python-MNE (Gramfort et al., 2013).

#### Pupil size

Pupil data was downsampled to 100 Hz and blinks were interpolated, with a cubic-spline interpolation or a linear interpolation if cubic-spline interpolation was not possible (Mathôt & Vilotijević, 2022). No baseline correction was applied to pupil size. The signal was cropped to the last 500 ms prior to stimulus presentation on each trial and the average was computed. Average pupil size was converted from arbitrary units to millimetres with a custom formula developed for the conditions of the lab. Trials with a physically impossible (< 2 mm or > 8 mm) pupil size were removed. For the main analyses, average pupil sizes were z-scored within participants and conditions, and trials that had a z-score three standard deviations away from the mean were removed.

#### Eye movements

To control for unwanted eye movements, we removed trials where gaze position significantly deviated from fixation. Specifically, trials where gaze position exceeded 4.6 dva from the centre for a minimum duration of 100ms were removed.

#### EEG

As a simple data quality check, topomaps and power spectral density plots were created based on the raw data. When these looked healthy based on subjective visual inspection (as was the case for all datasets), we proceeded with automatic processing of the EEG data as applied by the eeg-eyetracking-parser (Mathôt et al., 2023). During pre-processing data was re-referenced to the mastoids, the data was filtered with a 0.1 Hz high-pass filter, downsampled to 100 Hz, and an independent component analysis was applied to identify and remove blink-related artifacts. After pre-processing the data was epoched such that each epoch corresponded to the last 500 ms prior to stimulus presentation. When extracting epochs, the Autoreject algorithm (Jas et al., 2017) was applied to detect and interpolate bad channels and reject other artifacts (e.g., muscle movements).

##### Power estimation

To obtain average power in each band of interest in the pre-stimulus interval we first computed the PSD of the signal in the epoch spanning from 500 ms before to the moment of stimulus onset. We included posterior, parietal, central and temporal channels (O1, O2, Oz, POz, Pz, P3, P4, P7, P8, T7, T8, C3, Cz, C4). The PSD was computed using the MNE (Gramfort et al., 2013) function compute_psd with the default spectral estimation method (multitaper, using DPSS tapers (Slepian, 1978)). For the analysis, power in the theta (4–8 Hz), alpha (individually determined, as described below) and beta (12–30 Hz) bands was computed from the z-scored PSD. Specifically, z-scoring was applied per frequency bin and participant separately, by subtracting the mean and dividing by the standard deviation of that frequency bin across all trials and channels. The z-scored PSD was then averaged within each frequency band to obtain a trial-wise power estimate. Trials for which the resulting value was more than three standard deviations away from the participant’s mean were removed.

Individual alpha frequencies (IAF) were determined with the same procedure as in Samaha & Postle (2015), whereby the *electrode* with maximal power in the 8-13 Hz range was identified, and the *frequency* with maximal power in that electrode was considered the IAF peak. This procedure was chosen since most participants did not show a clear peak in the alpha range in the epoch of interest, presumably because alpha peaks are more prominent during a resting state than during task performance. The subject-specific electrodes and IAF peaks can be found in the Supplementary Materials. For the analysis we extracted average power in a range of ±1 Hz around the IAF peak.

##### 1/f slope computation

To complement the band-power analysis we also estimated the aperiodic component (1/f slope) of the power spectrum by fitting a linear regression to the log-log transformed PSD in the 6-30 Hz frequency range in the 500 ms pre-stimulus epoch. A more negative slope indicates more low-frequency power in the PSD. For analysis, the slopes were z-scored by subject and condition, and trials that had a z-score three standard deviations away from the mean were removed.

#### Skin conductance

The skin conductance signal was re-scaled from siemens (S) to microsiemens (μS) by multiplying the raw values by 1 000 000. The signal was then high-pass filtered with a 0.1 Hz filter. As for pupil and EEG measures, the average value in the 500 ms pre-stimulus epoch was computed and then subsequently z-scored by subject and condition. Trials that had a z-score three standard deviations away from the mean were removed.

## Analysis

We analysed the data using generalised linear mixed effects models constructed with the package lme4 in R (R Core Team, 2021). The main analysis consisted of three steps. First, we performed a model comparison procedure to determine which of the six variables may be relevant for explaining behaviour (accuracy and response tendency) on the task. The details of the procedure can be found below. Next, based on both the model comparison results and theoretical consideration we constructed another model including only a subset of the variables. Importantly, although the analysis pipeline was applied separately for the visual and auditory conditions and the model comparison results did not point to the same variables for each, we chose to include the same subset of variables in the final models for both conditions. This was done to allow for meaningful comparison between conditions. Finally, using the same subset of variables, we conducted a signal detection analysis on binned data to examine the influence of each variable on sensitivity (d’) and the criterion.

### Model comparison

We examined two different outcome variables and six possible explanatory variables. The outcome variables were accuracy and response, with response referring to whether participants reported detecting a stimulus regardless of whether it was actually present. The possible explanatory variables were target presence, pupil size, alpha power, beta power, theta power, 1/f slope, and skin conductance. We tested all possible combinations of the explanatory variables as main effects, but restricted interactions to those between target presence and each of the other explanatory variables. This reduced the number of tested models and total processing time. For the same reason, the random effects structure was limited to random intercepts for subjects. To identify relevant explanatory variables, we reviewed the ten best fitting models. Model fit was quantified with the Bayesian Information Criterion (BIC). For the sake of simplicity, we will not report the model comparison results in detail.

#### Main analysis

The variables that were significantly (*p <* 0.05) related to either accuracy or response in one of the experimental conditions were target presence, pupil size, and skin conductance. No interactions were significant. Therefore, these three variables were used as fixed effects for all main analyses. Since the random effects structure was restricted to only intercepts in the initial model comparison procedure, here we also tested whether model fit was improved by adding random slopes for each main effect. This was the case for all models.

#### Signal-detection analysis

The main analyses (described above) are conducted on individual trials, whereas signal-detection analysis is generally done on aggregate data. Although there have been studies attempting to conduct signal-detection analyses on single-trial data (Pilipenko & Samaha, 2024), in our hands this produced unstable outcomes. Therefore, we complement the main, single-trial analyses with traditional signal-detection analyses based on aggregate data.

To analyse the effects of pupil size and skin conductance on sensitivity (d’) and criterion, we first binned the data into 5-10 equally sized bins per subject, condition, and predictor, and then computed d’ and criterion in each bin. We then fitted a mixed linear model predicting d’ or criterion from the mean predictor value per bin, essentially treating the bin means as a continuous predictor. The model was fit with by-subject random intercepts. This was done separately for each combination of condition (auditory, visual), predictor (pupil and skin conductance), and dependent variable (d’, criterion). The entire analysis was repeated across bin counts 5-10 for robustness. In the results section we will report slope coefficients averaged across bin counts, with ranges of associated p-values.

## Results

### Behavioral performance

Behavioural performance measures (accuracy, hit rate, false alarm rate, sensitivity [d’], and criterion) are presented in Table 1, for each condition separately. The average accuracy, hit rate, false alarm rate, d’, and criterion for each individual subject can be found in the Supplementary Materials. Note that the statistics reported in Table 1 are computed on the subset of trials retained after removing bad trials based on data quality, as described above. Because this trial-level exclusion may shift per-subject accuracy relative to its original value, the accuracy of some participants on the retained trials exceeds the 90% cutoff used during screening, which was based on performance across all experimental trials.

**Table 1.**
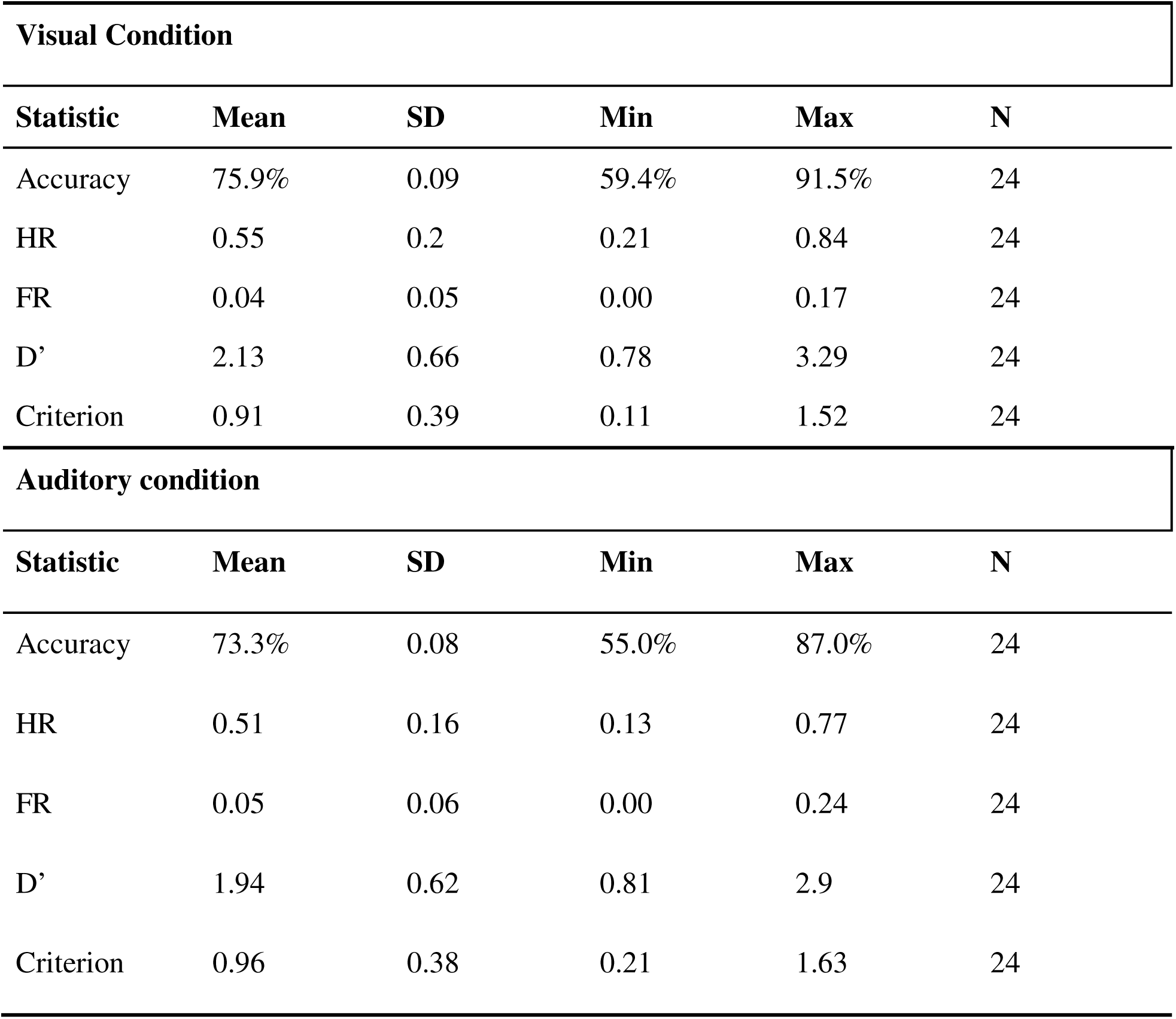
Behavioral performance.

### Time-on-task effects

Figure 2a depicts the trial-wise average pupil size (in mm), separately for the auditory and visual conditions. The dashed vertical lines indicate breaks in the task. The pupil is characteristically larger at the beginning of a block, constricting rapidly over the first few trials (Fig 2a). We did not correct for this pattern in the main analysis, as it reflects genuine fluctuations in pupil size. However, we conducted an additional analysis to control for potential effects of this systematic variation. Specifically, we fit a second-order polynomial with trial number as the predictor and pupil size as the dependent variable for each block separately. We then used the residual values and repeated the main analysis (cf. Claeys et al., 2026). This did not meaningfully change the pattern of results and we therefore conclude that the results are not driven by time-on-task. The details of the analysis and results are reported in the Supplementary Materials.

**Figure 2.**
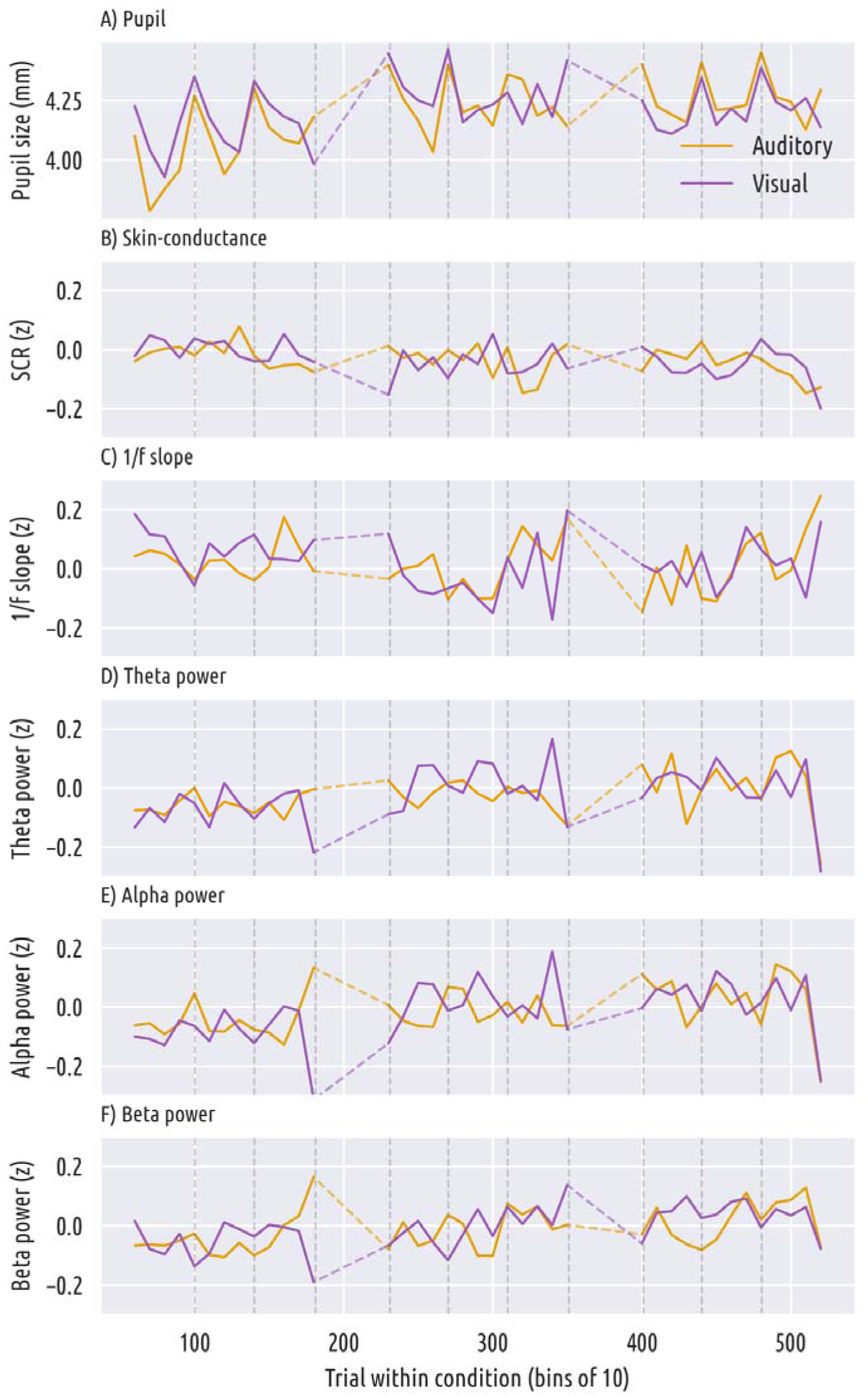
A) Variation in pupil size (in mm) over the course of the experiment in the visual (purple line) and auditory (orange line) conditions. Vertical dashed lines indicate breaks in the task, while horizontal dashes lines indicate staircase blocks which are removed from the analysis. To improve readability of the visualisation, pupil sizes are binned in bins of 10 trials for plotting. B) Variation in skin-conductance (z-scored) over the course of the experiment. C) Variation in 1/f slope (z-scored) over the course of the experiment. D) Variation in theta power (z-scored) over the course of the experiment. E) Variation in alpha power (z-scored) over the course of the experiment. F) Variation in beta power (z-scored) over the course of the experiment.

Skin conductance, as depicted in Figure 2b (z-scored) did not show systematic variation within individual blocks.

Figures 2c-f show the trial-wise variation in 1/f slope and EEG band power over time. None of the EEG measures show systematic variation in either condition. For further depiction of the EEG data topoplots, PSD plots and the Time-Frequency Spectrum of the pre-stimulus interval can be found in the Supplementary Materials.

### Average pupil size does not differ between conditions

Figure 3 shows the overall distribution and the per-subject average pupil sizes in each condition, measured in mm. The average pupil sizes were 4.22 mm and 4.18 mm in the visual and auditory conditions, respectively. To test whether the average size differed between conditions, we performed a paired-samples t-test, which indicated no significant differences (*t* = -0.59, *p* = 0.562).

**Figure 3.**
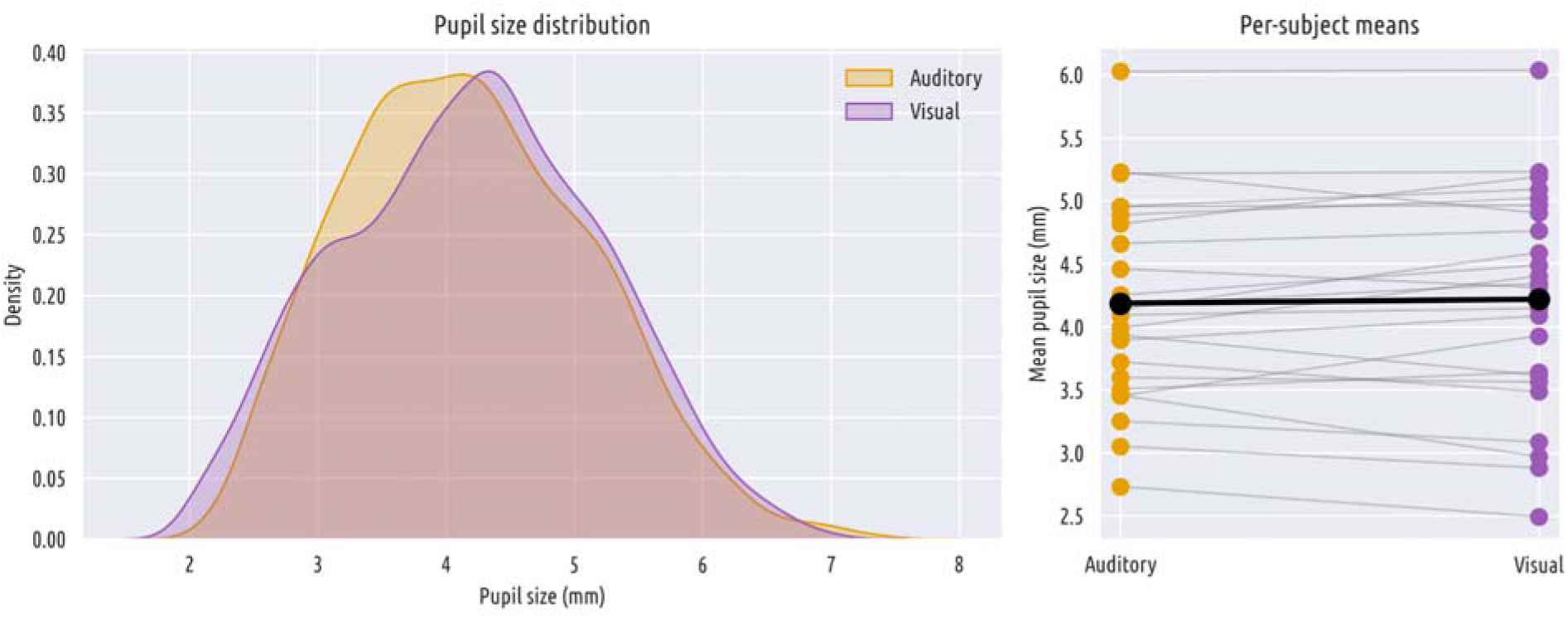
The distribution of recorded pupil sizes (on the left; in mm) and the average of each subject (on the right) in the visual (purple) and auditory (orange) conditions. Both conditions exhibit approximately normally distributed values, with no significant difference in the overall mean. Individual subjects differ in their average pupil size in general but there do not appear to be systematic differences between conditions.

### Relationships between explanatory variables

To examine the trial-level relationships between predictors, we first computed within-subject Pearson correlations (*r*) between each pair of predictors. Because r-values are not normally distributed, they should not be directly averaged. Therefore, the resulting subject-wise correlations were then transformed (with the Fisher z-transformation) and the transformed values were averaged. The resulting average was then back-transformed to obtain the correlation in the sample. These correlations are reported in Tables 2 and 3 for the visual and auditory conditions, respectively. The significance of the correlation was determined with a one-sample t-test on the transformed values of *r.* Relationships between pupil size and all other predictors are depicted in Figure 4. A full depiction of correlations between each pair of variables can be found in the Supplementary Materials.

**Figure 4.**
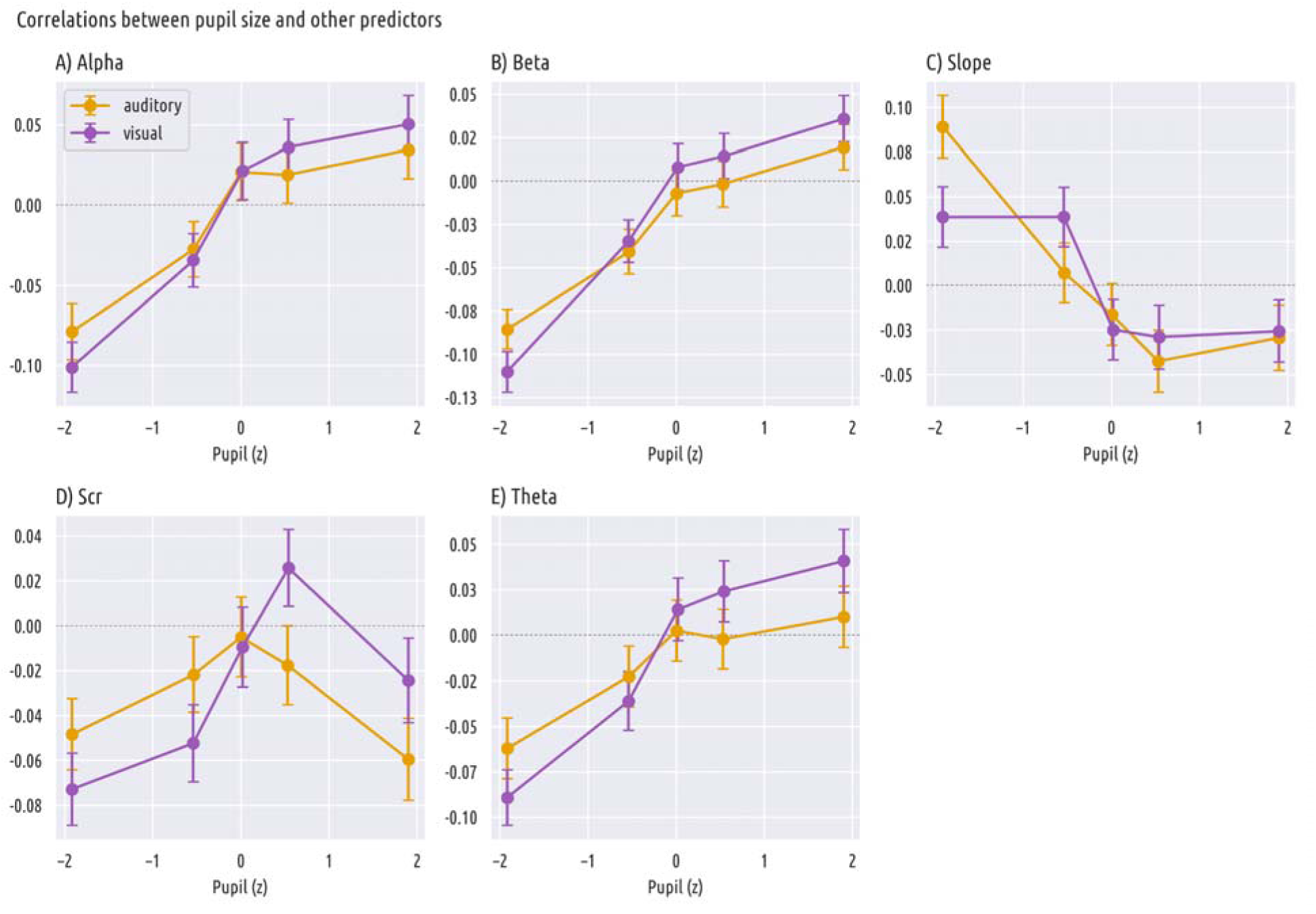
The average within-subject correlations between pupil size and other explanatory variables in the visual (purple) and auditory (orange) conditions. All values are normalized. Error bars represent the standard error of the mean (SEM) within the bin.

**Table 2.** Correlations between predictors in the visual condition.

|  | 1. | 2. | 3. | 4. | 5. | 6. |
| --- | --- | --- | --- | --- | --- | --- |
| 1. Pupil size | 1 |  |  |  |  |  |
| 2. Skin conductance | .03* | 1 |  |  |  |  |
| 3. Theta power | .06** | .02 | 1 |  |  |  |
| 4. Alpha power | .07** | .03 | .96*** | 1 |  |  |
| 5. Beta power | .1*** | .01 | .4*** | .51*** | 1 |  |
| 6. 1/f slope | -.02 | -.00 | -.63*** | -.65*** | .13** | 1 |

*Note:* \* $p < .05$ \*\* $p < .01$ \*\*\* $p < .001$

**Table 3.**
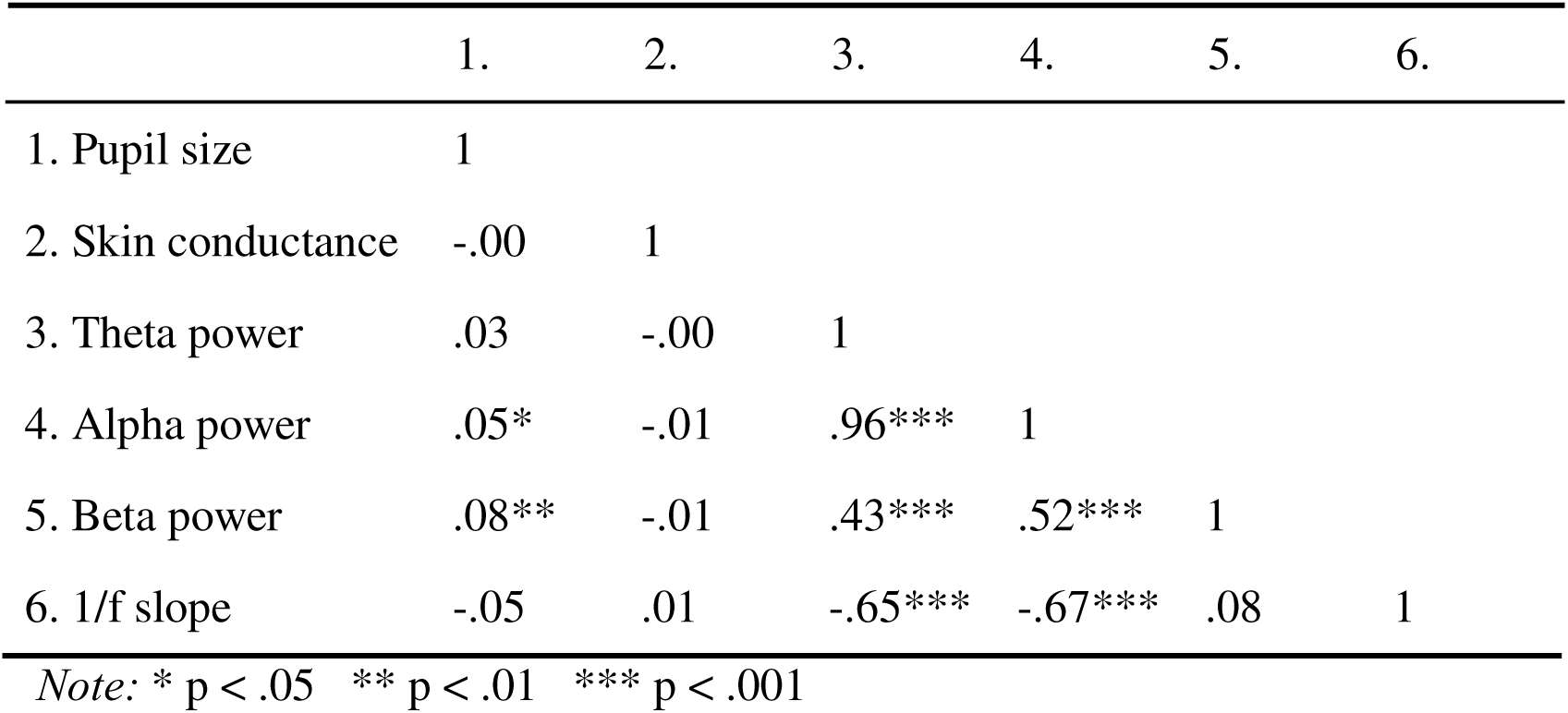
Correlations between predictors in the auditory condition.

|  | 1. | 2. | 3. | 4. | 5. | 6. |
| --- | --- | --- | --- | --- | --- | --- |
| 1. Pupil size | 1 |  |  |  |  |  |
| 2. Skin conductance | -.00 | 1 |  |  |  |  |
| 3. Theta power | .03 | -.00 | 1 |  |  |  |
| 4. Alpha power | .05* | -.01 | .96*** | 1 |  |  |
| 5. Beta power | .08** | -.01 | .43*** | .52*** | 1 |  |
| 6. 1/f slope | -.05 | .01 | -.65*** | -.67*** | .08 | 1 |

Note: \* $p < .05$ \*\* $p < .01$ \*\*\* $p < .001$

### Main analysis

The results of the main analyses are presented in figures 5 and 6, and reported in detail below. Figure 5 displays the model predictions obtained from the final fitted GLMs for pupil size and skin-conductance. Given that the fitted models were linear these prediction curves are also linear. To give a better image of the true shape of the relationship, figure 6 depicts the relationship between recorded pupil size and each outcome variable. For this plot, pupil size was binned into 5 equally sized bins, and the average accuracy, proportion of “yes”-responses, d’ and criterion in each bin was computed.

**Figure 5.**
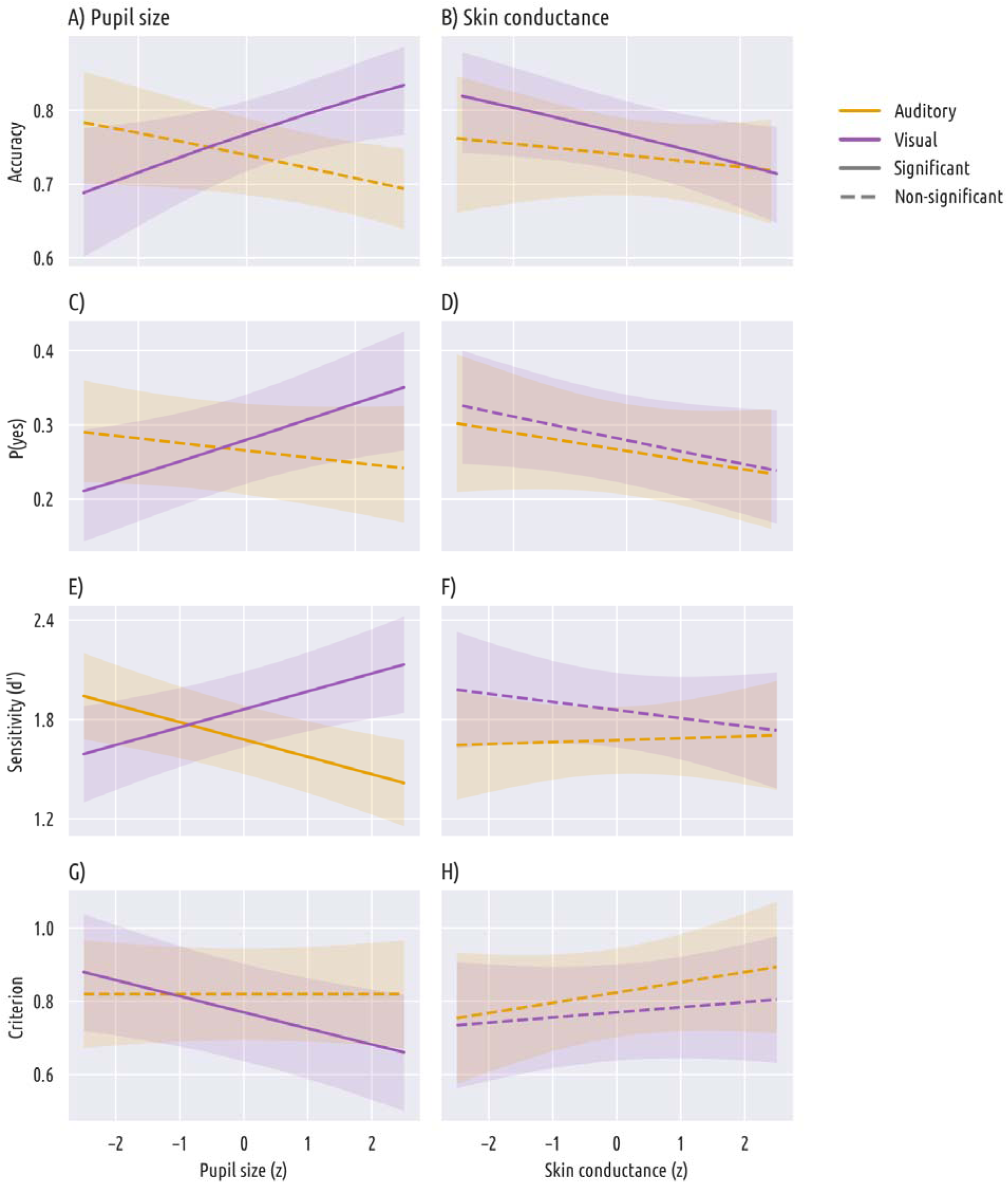
Overview of the results of the main analysis. The left column depicts the predicted effect of pupil size on each dependent variable (accuracy, response, sensitivity and criterion) and the right column depicts the predicted effect of SCR on each dependent variable. Predictions are derived from (generalized) linear mixed effects models. Purple lines represent the visual condition and orange lines the auditory condition. Solid lines indicate significant results and dashed lines indicate non-significant results.

**Figure 6.**
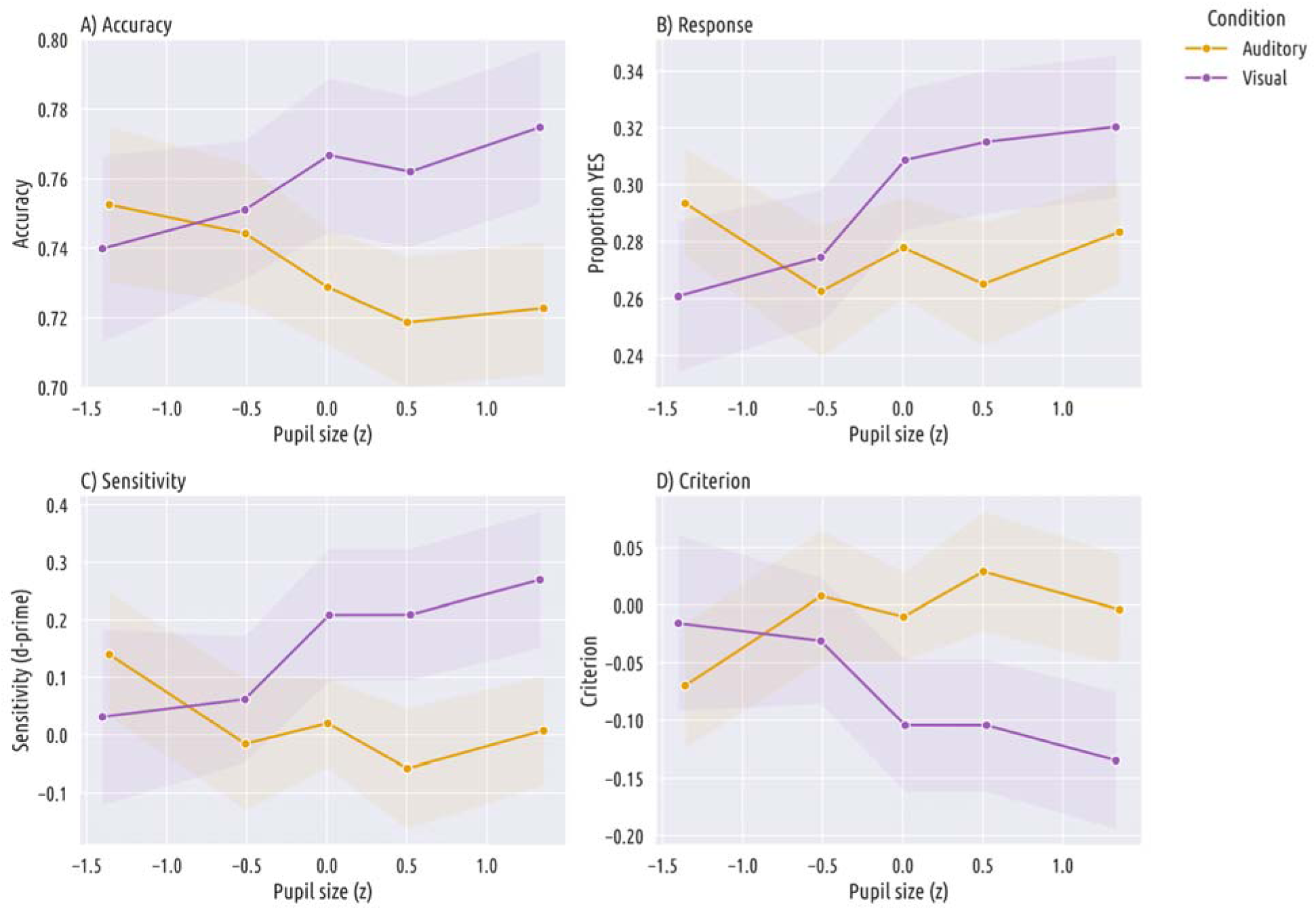
Each dependent variable as a function of binned pupil size. For visualization, z-scored pupil size was binned into 5 equally sized bins, and the average of each variable in the bin was calculated. Purple lines represent the visual condition and orange lines represent the auditory condition. A) Accuracy as a function of pupil size. B) Proportion of “yes”-responses as a function of pupil size. C) Sensitivity (d’) as a function of pupil size. D) Criterion as a function of pupil size.

### Pupil size and skin conductance are associated with accuracy in the visual condition

In the visual condition with accuracy as the dependent variable, there were significant effects of pupil size, target presence, and skin conductance. Larger pupils were associated with improved accuracy (*b* = 0.153, *p* = .02; Fig. 5a), while both the presence of a target (*b* = -3.683, *p* < .001) and higher skin conductance (*b* = -0.113, *p* = .049; Fig. 5b) were associated with decreased accuracy. (A negative effect of target presence on accuracy reflects that participants are more likely to miss targets that are actually present than they are to falsely report a target when it was not actually present.)

With response as the dependent variable, there were significant effects of pupil size and target presence. Both larger pupils (*b* = 0.137, *p* = .02; Fig. 5c) and the presence of a target (*b* = 4.194, *p* < .001) were associated with more ‘yes’-responses. Skin conductance was not significantly related to response tendency (*b* = -0.087, *p* = .113; Fig. 5d).

### Pupil size is not associated with differences in auditory detection performance

In the auditory condition with accuracy as the dependent variable, there was a significant effect of target presence, such that the presence of a target was associated with decreased accuracy (*b* = -3.823, *p* < .001). The effects of pupil size (*b* = -0.09, *p* = .053; Fig. 5a) and skin conductance (*b* = -0.048, *p* = .498; Fig. 5b) were not significant.

With response as the dependent variable, there was a significant effect of target presence, such that the presence of a target was associated with more ‘yes’-responses (*b* = 3.887, *p* < .001). Pupil size (*b* = -0.048, *p* = .274; Fig. 5c) and skin conductance (*b* = -0.073, *p* = .297; Fig. 5d) were not significantly related to response tendency.

### Larger pupils are associated with higher sensitivity and a more liberal criterion in visual detection

In the visual condition the SDT analysis suggested that larger pupils are associated with higher sensitivity (Fig. 5e) and a more liberal criterion (Fig. 5g). The effect of pupil size on d’ reached significance with all bin counts. The average slope coefficient was *b* = 0.10 and the p-value ranged from *p* = .001 to *p* = .01. The effect of pupil size on criterion also reached significance with all bin counts. The average slope coefficient was -0.05 and the p-value ranged from *p* = .004 to *p* = .03.

There were no significant effects of skin conductance on either d’ (Fig. 5f) or criterion (Fig. 5h).

### Larger pupils are associated with lower sensitivity in auditory detection

In the auditory condition the SDT analysis suggested that larger pupils are associated with lower sensitivity. The effect of pupil size on d’ reached significance with all bin counts. The average slope coefficient was *b* = -0.10 and the p-value ranged from *p* = .001 to *p* = .01. There were no significant effects on the criterion.

There were no significant effects of skin conductance on either d’ or criterion.

## Discussion

Here, we investigated the effects of pupil size, skin conductance, and EEG power variation on detection performance in the visual and auditory modalities. Our aim was to better understand the role of spontaneous pupil-size fluctuations in perceptual processing. Previous research suggested that the optical consequences of pupil size changes affect visual detection performance independently of fluctuations in arousal (Ruuskanen et al., 2025; Ruuskanen & Mathôt, 2026a). However, studies have so far not been able to rigorously dissociate optical- and arousal-related mechanisms. This is what we set out to do here.

Given this aim, the most important aspects of our results are related to the dissociation between optics and arousal. The results that most directly support this dissociation are the opposite effect of pupil size on performance in the visual and auditory conditions, and the opposite effects of pupil size and skin-conductance on performance in the visual condition. Firstly, in the visual condition we found that larger pupils were associated with higher accuracy and more target-present responses, replicating previous research (Eberhardt et al., 2022; Mathôt & Ivanov, 2019; Ruuskanen et al., 2025; Ruuskanen & Mathôt, 2026b). Furthermore, and also consistent with previous findings, signal-detection analyses showed that pupil size was robustly associated with improved sensitivity and a more liberal criterion (Beerendonk et al., 2024; de Gee et al., 2014; Podvalny et al., 2021). Conversely, in the auditory condition, where any relationship between pupil size and performance is assumed to be driven solely by arousal, larger pupils were associated with lower performance, reflected especially in lower sensitivity. We suggest that these opposite effects reflect the presence of an optical effect in the visual condition, which drives an overall large-pupil advantage for visual detection by allowing more light to enter the eye (thus improving sensitivity by increasing signal-to-noise ratio). This conclusion is further supported by the finding that in the visual condition another measure related to arousal–skin-conductance–had the opposite effect on performance from pupil size, whereby higher skin-conductance was associated with lower accuracy. This was despite pupil size and skin-conductance being positively correlated with one another (Chang et al., 2025). Based on the diverging results between conditions and measures, we conclude that in visual detection the relationship between pupil size and performance is driven by a combination of optics and arousal (Ruuskanen & Mathôt, 2026a), whereas in the auditory condition the relationship is driven purely by arousal.

Additional evidence for the optical account comes from the shape of the pupil-performance relationship in the visual condition, which is inconsistent with what a purely arousal-based account would predict. Specifically, whereas sensitivity is typically expected to show an inverted-U relationship with pupil size (peaking at intermediate pupil size) (Beerendonk et al., 2024; Podvalny et al., 2021) we observe a rather linear relationship, with the largest pupils associated with the highest sensitivity; the same holds for accuracy. This further supports our conclusion that the relationship is not solely driven by arousal, as a purely arousal-driven account would predict an inverted-U.

In addition to these relatively straightforward results, we uncovered several other patterns that are more difficult to interpret, with the first being the relationship between pupil size and performance in the auditory condition. With some exceptions (Doll et al., 2024) it is generally found that performance in auditory and tactile tasks tends to display an inverted-U shaped relationship with pupil size (Beerendonk et al., 2024; Nuiten et al., 2026). Accordingly, this is also what we expected to observe. However, our results deviate from this expectation. The relationship between pupil size and sensitivity in the auditory condition displayed an almost exponential decrease, with lower sensitivity for all but the smallest pupil sizes. Using accuracy as the outcome measure also revealed a negative relationship, though less extreme and with a slight increase at medium pupil sizes. While this pattern is somewhat more in line with our initial expectations, it still does not resemble an inverted-U shape, and the effect on accuracy was less robust than on sensitivity. One possible explanation for this pattern concerns the measured range of arousal. It has been suggested that differences across studies may reflect sampling of different portions of the arousal spectrum: studies showing linear increases may predominantly sample the lower range, while those showing the full inverted-U sample a broader range (Beerendonk et al., 2024). Following this logic, our auditory condition may have sampled the upper range of the arousal spectrum — perhaps because participants perceived the auditory task as more difficult, pushing them into a higher arousal state. However, the ranges and averages of measured pupil sizes did not differ significantly between conditions, suggesting that baseline arousal and perceived effort were similar across the visual and auditory tasks. Why auditory performance deteriorates at larger pupil sizes therefore remains unclear.

Similarly, against initial expectations, neither EEG power nor the aperiodic component (1/f slope) were related to performance differences on the task. This is somewhat puzzling, given a large literature on the relationship between alpha power and sensory processing, where alpha suppression is generally associated with criterion-changes (Iemi et al., 2017; Limbach & Corballis, 2016; Samaha et al., 2017, 2020), and with increased hit-rate or accuracy (Ergenoglu et al., 2004; Melcón et al., 2024). However, null results have been reported before (Ruuskanen et al., 2025) and some authors argue that alpha phase may be more important than average power in determining performance (Busch et al., 2009; Pilipenko et al., 2026). We also failed to replicate previously reported links between performance and power in the theta and beta bands (Podvalny et al., 2021; Ruuskanen et al., 2025).

Importantly, we did replicate the previously observed positive correlation between pupil size and alpha and beta band power (Mathôt et al., 2023; Ruuskanen et al., 2025) suggesting that there was sufficient variation in EEG power for performance differences to emerge had they existed. However, the positive correlation between pupil size and broadband EEG power is itself a somewhat puzzling finding. During a resting state, a correlation between pupil size and alpha power is generally observed (Montefusco-Siegmund et al., 2022), but during task performance it is more variable and not always replicated (Ruuskanen & Mathôt, 2026a). Given that both alpha suppression and pupil dilation are commonly assumed to reflect increased arousal, a negative correlation between the two would be expected; yet a positive one is typically found. One complicating factor is that alpha power is thought to reflect not only arousal but also cortical excitability (Lange et al., 2013; Romei et al., 2008), and while the two are related, they are not equivalent (Barry et al., 2020; Schubring & Schupp, 2021; Weiss et al., 2026). Given this complexity, the origin of the positive pupil-alpha correlation remains an open question. On the other hand, the positive correlation between beta power and pupil size has received less attention, but seems to be more robust (Mathôt et al., 2023; Ruuskanen et al., 2025; Waschke et al., 2019). Similarly to alpha power, lower beta power has been related to increased arousal (Schubring & Schupp, 2021), thus presenting the same conundrum: why is broadband EEG power positively correlated with pupil size? In the case of beta power the answer may lie in motor control, which beta power is associated with (Engel & Fries, 2010; Jenkinson & Brown, 2011). The positive correlation between pupil size and power in the beta band may be a reflection of motor control of the iris dilator muscle.

In broader terms, our results have implications for the important questions of what different arousal measures capture, and how arousal relates to perceptual performance. A key observation in this regard is that while pupil size was positively related to detection performance and skin conductance was negatively related to detection performance, the two measures were themselves positively correlated with each other. This pattern is consistent with the proposed multidimensional nature of arousal (Sabat et al., 2025): the positive correlation between the two measures suggests they share a domain-general arousal component, while their opposite effects on performance suggest they also tap into distinct arousal domains. A tentative proposal is that pupil size reflects a more cognitive form of arousal, closely tied to neuromodulatory systems such as the locus coeruleus-norepinephrine system, while skin conductance more strongly reflects physiological arousal. The lack of performance-related effects of EEG power, despite its correlation with pupil size, further emphasises that different arousal measures are not interchangeable. Future research should aim to better characterize the distinct contributions of different arousal domains to sensory processing and perceptual decision-making.

To recap, our findings support the hypothesis that in visual detection tasks the relationship between pupil size and performance is driven by two underlying mechanisms: arousal and optics (Ruuskanen & Mathôt, 2026a). More generally, we suggest that the optical consequences of pupil size in visual detection represent a subtle form of sensory tuning, which refers to the adjustment of the visual system to current task demands (Mathôt, 2020). This also provides a tentative answer to the often overlooked fundamental question: *why* does pupil size increase with arousal? We suggest sensory tuning as the functional explanation. Highly arousing situations are generally also associated with a need for vigilance, where visual sensitivity (provided by large pupils) is more important than visual acuity (associated with small pupils; for more in-depth discussion see, Ruuskanen & Mathôt, 2026a). From this perspective, pupil dilation in response to arousal is not merely a physiological byproduct, but an adaptive mechanism that prepares the visual system for the perceptual demands of high-arousal situations.

In conclusion, larger pupils were associated with higher sensitivity in visual detection and lower sensitivity in auditory detection. EEG measures, including broadband power and the aperiodic 1/f component, were not related to performance, though we did replicate the previously observed positive correlations between pupil size and alpha and beta band power. Taken together, these results support the view that the effect of pupil size on visual detection performance is driven by two distinct mechanisms: arousal and optics, while in auditory detection the relationship is solely driven by arousal. More broadly, our findings highlight the pupil as an active and functional component of visual processing, rather than simply a measure of arousal.

## References

Barry, R. J., De Blasio, F. M., Fogarty, J. S., & Clarke, A. R. (2020). Natural alpha frequency components in resting EEG and their relation to arousal. Clinical Neurophysiology, 131(1), 205–212. 10.1016/j.clinph.2019.10.018

Beerendonk, L., Mejías, J. F., Nuiten, S. A., de Gee, J. W., Fahrenfort, J. J., & van Gaal, S. (2024). A disinhibitory circuit mechanism explains a general principle of peak performance during mid-level arousal. Proceedings of the National Academy of Sciences, 121(5), e2312898121. 10.1073/pnas.2312898121

Boncompte, G., Villena-González, M., Cosmelli, D., & López, V. (2016). Spontaneous Alpha Power Lateralization Predicts Detection Performance in an Un-Cued Signal Detection Task. PLOS ONE, 11(8), e0160347. 10.1371/journal.pone.0160347

Busch, N. A., Dubois, J., & VanRullen, R. (2009). The Phase of Ongoing EEG Oscillations Predicts Visual Perception. Journal of Neuroscience, 29(24), 7869–7876. 10.1523/JNEUROSCI.0113-09.2009

Chang, Y.-H., Yep, R., & Wang, C.-A. (2025). Pupil size correlates with heart rate, skin conductance, pulse wave amplitude, and respiration responses during emotional conflict and valence processing. Psychophysiology, 62(1), e14726. 10.1111/psyp.14726

Claeys, W., Ruuskanen, V., & Mathot, S. (2026). Subtle Pupil-Size Changes Associated With Exploration Do Not Affect Visual Sensitivity (p. 2026.06.05.730459). bioRxiv. 10.64898/2026.06.05.730459

Dalmaijer, E. S., Mathôt, S., & Van der Stigchel, S. (2014). PyGaze: An open-source, cross-platform toolbox for minimal-effort programming of eyetracking experiments. Behavior Research Methods, 46(4), 913–921. 10.3758/s13428-013-0422-2

de Gee, J. W., Knapen, T., & Donner, T. H. (2014). Decision-related pupil dilation reflects upcoming choice and individual bias. Proceedings of the National Academy of Sciences, 111(5), E618–E625. 10.1073/pnas.1317557111

de Gee, J. W., Mridha, Z., Hudson, M., Shi, Y., Ramsaywak, H., Smith, S., Karediya, N., Thompson, M., Jaspe, K., Jiang, H., Zhang, W., & McGinley, M. J. (2024). Strategic stabilization of arousal boosts sustained attention. Current Biology, 34(18), 4114–4128.e6. 10.1016/j.cub.2024.07.070

Doll, L., Dykstra, A. R., & Gutschalk, A. (2024). Perceptual awareness of near-threshold tones scales gradually with auditory cortex activity and pupil dilation. iScience, 27(8). 10.1016/j.isci.2024.110530

Doll, L., Heiland, S., & Gutschalk, A. (2025). A Role of Pupil-linked Arousal, Cingulo-insular Cortex, and Intralaminar Thalamus for Auditory Near-threshold Perception. Journal of Cognitive Neuroscience, 1–25. 10.1162/jocn_a_02324

Eberhardt, L. V., Strauch, C., Hartmann, T. S., & Huckauf, A. (2022). Increasing pupil size is associated with improved detection performance in the periphery. *Attention, Perception*, & Psychophysics, 84(1), 138–149. 10.3758/s13414-021-02388-w

Engel, A. K., & Fries, P. (2010). Beta-band oscillations—Signalling the status quo? Current Opinion in Neurobiology, 20(2), 156–165. 10.1016/j.conb.2010.02.015

Ergenoglu, T., Demiralp, T., Bayraktaroglu, Z., Ergen, M., Beydagi, H., & Uresin, Y. (2004). Alpha rhythm of the EEG modulates visual detection performance in humans. Cognitive Brain Research, 20(3), 376–383. 10.1016/j.cogbrainres.2004.03.009

Gramfort, A., Luessi, M., Larson, E., Engemann, D. A., Strohmeier, D., Brodbeck, C., Goj, R., Jas, M., Brooks, T., Parkkonen, L., & Hämäläinen, M. (2013). MEG and EEG data analysis with MNE-Python. Frontiers in Neuroscience, 7. 10.3389/fnins.2013.00267

Grujic, N., Polania, R., & Burdakov, D. (2024). Neurobehavioral meaning of pupil size. Neuron, 112(20), 3381–3395. 10.1016/j.neuron.2024.05.029

Iemi, L., Chaumon, M., Crouzet, S. M., & Busch, N. A. (2017). Spontaneous Neural Oscillations Bias Perception by Modulating Baseline Excitability. Journal of Neuroscience, 37(4), 807–819. 10.1523/JNEUROSCI.1432-16.2016

Jas, M., Engemann, D. A., Bekhti, Y., Raimondo, F., & Gramfort, A. (2017). Autoreject: Automated artifact rejection for MEG and EEG data. NeuroImage, 159, 417–429. 10.1016/j.neuroimage.2017.06.030

Jenkinson, N., & Brown, P. (2011). New insights into the relationship between dopamine, beta oscillations and motor function. Trends in Neurosciences, 34(12), 611–618. 10.1016/j.tins.2011.09.003

Joshi, S., Li, Y., Kalwani, R. M., & Gold, J. I. (2016). Relationships between Pupil Diameter and Neuronal Activity in the Locus Coeruleus, Colliculi, and Cingulate Cortex. Neuron, 89(1), 221–234. 10.1016/j.neuron.2015.11.028

Koenig, L., & He, B. J. (2025). Spontaneous slow cortical potentials and brain oscillations independently influence conscious visual perception. PLOS Biology, 23(1), e3002964. 10.1371/journal.pbio.3002964

Lange, J., Oostenveld, R., & Fries, P. (2013). Reduced Occipital Alpha Power Indexes Enhanced Excitability Rather than Improved Visual Perception. Journal of Neuroscience, 33(7), 3212–3220. 10.1523/JNEUROSCI.3755-12.2013

Lendner, J. D., Helfrich, R. F., Mander, B. A., Romundstad, L., Lin, J. J., Walker, M. P., Larsson, P. G., & Knight, R. T. (2020). An electrophysiological marker of arousal level in humans. eLife, 9, e55092. 10.7554/eLife.55092

Limbach, K., & Corballis, P. M. (2016). Prestimulus alpha power influences response criterion in a detection task. Psychophysiology, 53(8), 1154–1164. 10.1111/psyp.12666

Mathôt, S. (2020). Tuning the Senses: How the Pupil Shapes Vision at the Earliest Stage. Annual Review of Vision Science, 6, 433–451. 10.1146/annurev-vision-030320-062352

Mathôt, S., Berberyan, H., Büchel, P., Ruuskanen, V., Vilotijević, A., & Kruijne, W. (2023). Effects of pupil size as manipulated through ipRGC activation on visual processing. NeuroImage, 283, 120420. 10.1016/j.neuroimage.2023.120420

Mathôt, S., & Ivanov, Y. (2019). The effect of pupil size and peripheral brightness on detection and discrimination performance. PeerJ, 7, e8220. 10.7717/peerj.8220

Mathôt, S., Schreij, D., & Theeuwes, J. (2012). OpenSesame: An open-source, graphical experiment builder for the social sciences. Behavior Research Methods, 44(2), 314–324. 10.3758/s13428-011-0168-7

Mathôt, S., & Vilotijević, A. (2022). Methods in cognitive pupillometry: Design, preprocessing, and statistical analysis. Behavior Research Methods. 10.3758/s13428-022-01957-7

McGinley, M. J., David, S. V., & McCormick, D. A. (2015). Cortical Membrane Potential Signature of Optimal States for Sensory Signal Detection. Neuron, 87(1), 179–192. 10.1016/j.neuron.2015.05.038

Melcón, M., Stern, E., Kessel, D., Arana, L., Poch, C., Campo, P., & Capilla, A. (2024). Perception of near-threshold visual stimuli is influenced by prestimulus alpha-band amplitude but not by alpha phase. Psychophysiology, 61(5), e14525. 10.1111/psyp.14525

Montefusco-Siegmund, R., Schwalm, M., Rosales Jubal, E., Devia, C., Egaña, J. I., & Maldonado, P. E. (2022). Alpha EEG Activity and Pupil Diameter Coupling during Inactive Wakefulness in Humans. eNeuro, 9(2), ENEURO.0060-21.2022. 10.1523/ENEURO.0060-21.2022

Murphy, P. R., Robertson, I. H., Balsters, J. H., & O’connell, R. G. (2011). Pupillometry and P3 index the locus coeruleus-noradrenergic arousal function in humans. Psychophysiology, 48(11), 1532–1543. 10.1111/j.1469-8986.2011.01226.x

Nuiten, S. A., De Gee, J. W., Zantvoord, J. B., Sterzer, P., Fahrenfort, J. J., & van Gaal, S. (2026). Phasic and tonic arousal distinctly shape human decision bias. Communications Biology, 9(1), 553. 10.1038/s42003-026-09776-8

Peirce, J., Gray, J. R., Simpson, S., MacAskill, M., Höchenberger, R., Sogo, H., Kastman, E., & Lindeløv, J. K. (2019). PsychoPy2: Experiments in behavior made easy. Behavior Research Methods, 51(1), 195–203. 10.3758/s13428-018-01193-y

Pilipenko, A., Mcgowan, A., & Samaha, J. (2026). Alpha-Band Phase Modulates Perceptual Sensitivity by Changing Internal Noise and Sensory Tuning. eLife, 15. 10.7554/eLife.110000.2

Pilipenko, A., & Samaha, J. (2024). Double dissociation of spontaneous alpha-band activity and pupil-linked arousal on additive and multiplicative perceptual gain. Journal of Neuroscience. 10.1523/JNEUROSCI.1944-23.2024

Podvalny, E., King, L. E., & He, B. J. (2021). Spectral signature and behavioral consequence of spontaneous shifts of pupil-linked arousal in human. eLife, 10, e68265. 10.7554/eLife.68265

R Core Team. (2021). R: A language and environment for statistical computing. [Computer software]. R Foundation for Statistical Computing. https://www.R-project.org/

Reimer, J., McGinley, M. J., Liu, Y., Rodenkirch, C., Wang, Q., McCormick, D. A., & Tolias, A. S. (2016). Pupil fluctuations track rapid changes in adrenergic and cholinergic activity in cortex. Nature Communications, 7(1), 13289. 10.1038/ncomms13289

Renard, Y., Lotte, F., Gibert, G., Congedo, M., Maby, E., Delannoy, V., Bertrand, O., & Lécuyer, A. (2010). OpenViBE: An Open-Source Software Platform to Design, Test, and Use Brain–Computer Interfaces in Real and Virtual Environments. Presence: Teleoperators and Virtual Environments, 19(1), 35–53. 10.1162/pres.19.1.35

Romei, V., Brodbeck, V., Michel, C., Amedi, A., Pascual-Leone, A., & Thut, G. (2008). Spontaneous fluctuations in posterior alpha-band EEG activity reflect variability in excitability of human visual areas. Cerebral Cortex (New York, N.Y.: 1991), 18(9), 2010– 2018. 10.1093/cercor/bhm229

Ruuskanen, V., Boehler, C. N., & Mathôt, S. (2025). The Interplay of Spontaneous Pupil-Size Fluctuations and EEG Power in Near-Threshold Detection. Psychophysiology, 62(3), e70035. 10.1111/psyp.70035

Ruuskanen, V., & Mathôt, S. (2026a). *Beyond Arousal: Pupil Fluctuations, Neural Activity, and Behavior* (Mkdgc_v1). PsyArXiv. https://osf.io/preprints/psyarxiv/mkdgc_v1/

Ruuskanen, V., & Mathôt, S. (2026b). Pupil size correlates with near-threshold detection performance irrespective of stimulus color, eccentricity, or retinal adaptation state. Journal of Experimental Psychology: Human Perception and Performance. 10.1037/xhp0001417

Sabat, M., de Dampierre, C., & Tallon-Baudry, C. (2025). Evidence for domain-general arousal from semantic and neuroimaging meta-analyses reconciles opposing views on arousal. Proceedings of the National Academy of Sciences of the United States of America, 122(6), e2413808122. 10.1073/pnas.2413808122

Samaha, J., Iemi, L., Haegens, S., & Busch, N. A. (2020). Spontaneous Brain Oscillations and Perceptual Decision-Making. Trends in Cognitive Sciences, 24(8), 639–653. 10.1016/j.tics.2020.05.004

Samaha, J., Iemi, L., & Postle, B. R. (2017). Prestimulus alpha-band power biases visual discrimination confidence, but not accuracy. *Consciousness and Cognition*, Time Course of Event-Related Potentials Associated with Conscious Experience, 54, 47–55. 10.1016/j.concog.2017.02.005

Samaha, J., & Postle, B. R. (2015). The Speed of Alpha-Band Oscillations Predicts the Temporal Resolution of Visual Perception. Current Biology, 25(22), 2985–2990. 10.1016/j.cub.2015.10.007

Schriver, B. J., Bagdasarov, S., & Wang, Q. (2018). Pupil-linked arousal modulates behavior in rats performing a whisker deflection direction discrimination task. Journal of Neurophysiology. (Bethesda, MD). 10.1152/jn.00290.2018

Schubring, D., & Schupp, H. T. (2021). Emotion and Brain Oscillations: High Arousal is Associated with Decreases in Alpha- and Lower Beta-Band Power. Cerebral Cortex, 31(3), 1597–1608. 10.1093/cercor/bhaa312

Slepian, D. (1978). Prolate spheroidal wave functions, fourier analysis, and uncertainty — V: The discrete case. The Bell System Technical Journal, 57(5), 1371–1430. The Bell System Technical Journal. 10.1002/j.1538-7305.1978.tb02104.x

Vilotijević, A., & Mathôt, S. (2024). Functional benefits of cognitively driven pupil-size changes. WIREs Cognitive Science, 15(3), e1672. 10.1002/wcs.1672

Wang, C.-A., Baird, T., Huang, J., Coutinho, J. D., Brien, D. C., & Munoz, D. P. (2018). Arousal Effects on Pupil Size, Heart Rate, and Skin Conductance in an Emotional Face Task. Frontiers in Neurology, 9. 10.3389/fneur.2018.01029

Waschke, L., Tune, S., & Obleser, J. (2019). Local cortical desynchronization and pupil-linked arousal differentially shape brain states for optimal sensory performance. eLife, 8, e51501. 10.7554/eLife.51501

Watson, A. B., & Pelli, D. G. (1983). Quest: A Bayesian adaptive psychometric method. Perception & Psychophysics, 33(2), 113–120. 10.3758/BF03202828

Yerkes, R. M., & Dodson, J. D. (1908). The relation of strength of stimulus to rapidity of habit□formation. Journal of Comparative Neurology and Psychology, 18(5), 459–482. 10.1002/cne.920180503

