## Supplementary Materials for "Larger Pupils are Associated with Improved Visual, but not Auditory, Near-Threshold Detection"

### IAF peaks and electrodes

Table 1 shows for each subject (that was included in the analysis) the electrode with maximal power in the 8-13Hz range and the frequency with maximal power in that electrode.

| **Table 1**  *Subject-specific IAF electrodes and frequencies* | | |
| --- | --- | --- |
| Subject | Channel | Frequency |
| 2 | P3 | 8.96 |
| 3 | Pz | 8.96 |
| 4 | Pz | 8.96 |
| 5 | POz | 8.96 |
| 6 | POz | 9.96 |
| 7 | POz | 9.96 |
| 9 | POz | 9.96 |
| 14 | Pz | 8.96 |
| 15 | Pz | 8.96 |
| 16 | Pz | 8.96 |
| 17 | P3 | 9.96 |
| 18 | Pz | 12.95 |
| 19 | POz | 8.96 |
| 20 | POz | 9.96 |
| 21 | Pz | 8.96 |
| 20 | POz | 8.96 |
| 21 | POz | 8.96 |
| 22 | POz | 8.96 |
| 23 | P3 | 8.96 |
| 24 | Pz | 8.96 |
| 25 | POz | 8.96 |
| 26 | P3 | 8.96 |
| 27 | POz | 8.96 |
| 28 | P7 | 9.96 |
| 29 | P3 | 8.96 |
| 32 | P7 | 9.96 |
| 33 | P4 | 8.96 |

### Individual performance metrics

Table 2 has the average performance metrics (per condition) for each subject included in the analysis. Note that these are computed after the data was processed, meaning that they are based only on the trials included in the analysis.

| **Table 2**  *Subject-wise performance metrics: Auditory condition* | | | | | |
| --- | --- | --- | --- | --- | --- |
| Subject | Accuracy | Hit rate | FA rate | D’ | Criterion |
| 2 | .818 | .623 | .006 | 2.838 | 1.104 |
| 3 | .651 | .302 | .003 | 2.231 | 1.633 |
| 4 | .688 | .430 | .005 | 2.393 | 1.367 |
| 5 | .696 | .475 | .065 | .450 | .788 |
| 6 | .758 | .551 | .044 | 1.836 | .789 |
| 7 | .559 | .134 | .016 | 1.028 | 1.622 |
| 9 | .870 | .771 | .038 | 2.515 | .514 |
| 14 | .795 | .771 | .038 | 2.515 | .514 |
| 15 | .683 | .391 | .010 | 2.051 | 1.303 |
| 16 | .720 | .466 | .023 | 1.853 | 1.062 |
| 17 | .550 | .125 | .025 | .810 | 1.555 |
| 18 | .831 | .689 | .039 | 2.254 | .634 |
| 19 | .697 | .616 | .236 | 1.014 | .213 |
| 20 | .719 | .466 | .044 | 1.626 | 0.897 |
| 21 | .825 | .655 | .009 | 2.760 | .980 |
| 22 | .677 | .383 | .040 | 1.447 | 1.022 |
| 24 | .683 | .516 | .150 | 1.075 | .497 |
| 25 | .715 | .634 | .205 | 1.166 | .242 |
| 26 | .719 | .469 | .017 | 2.050 | 1.103 |
| 27 | .790 | .571 | .003 | 2.901 | 1.271 |
| 28 | .745 | .530 | .040 | 1.825 | .837 |
| 29 | .798 | .619 | .020 | 2.349 | .872 |
| 32 | .789 | .601 | .027 | 2.185 | .836 |
| 33 | .825 | .662 | .020 | 2.464 | .815 |
| *Subject-wise performance metrics: Visual condition* | | | | | |
| Subject | Accuracy | Hit rate | FA rate | D’ | Criterion |
| 2 | .640 | .319 | .005 | 2.084 | 1.512 |
| 3 | .685 | .376 | .016 | 1.830 | 1.231 |
| 4 | .655 | .332 | .014 | 1.769 | 1.319 |
| 5 | .674 | .347 | .018 | 1.710 | 1.248 |
| 6 | .789 | .637 | .065 | 1.865 | .582 |
| 7 | .706 | .399 | .011 | 2.025 | 1.268 |
| 9 | .731 | .488 | .038 | 1.744 | .902 |
| 14 | .596 | .212 | .025 | 1.161 | 1.379 |
| 15 | .601 | .293 | .093 | .778 | .933 |
| 16 | .778 | .565 | .004 | 2.854 | 1.263 |
| 17 | .673 | .383 | .027 | 1.634 | 1.114 |
| 18 | .896 | .795 | .009 | 3.198 | .776 |
| 19 | .828 | .722 | .085 | 1.963 | .392 |
| 20 | .890 | .792 | .020 | 2.863 | .618 |
| 21 | .890 | .814 | .038 | 2.668 | .440 |
| 22 | .734 | .457 | .010 | 2.219 | 1.217 |
| 24 | .770 | .713 | .173 | 1.505 | .191 |
| 25 | .805 | .772 | .165 | 1.718 | .113 |
| 26 | .745 | .532 | .034 | 1.910 | .876 |
| 27 | .915 | .844 | .018 | 3.107 | .544 |
| 28 | .766 | .533 | .003 | 2.831 | 1.333 |
| 29 | .723 | .480 | .032 | 1.805 | .952 |
| 32 | .821 | .665 | .015 | 2.605 | .876 |
| 33 | .908 | .825 | .010 | 3.274 | .702 |

### Results of time-on-task analysis

To rule out potential effects of systematic variation in pupil size due to time-on-task, whereby pupil size decreases within each block, we fitted a second-order polynomial pupil size within a block. We then took the residual values of the polynomial and repeated the main analysis, meaning fitted two GLM’s (one with only random intercepts and one with random slopes) on the data, with pupil size (residual), target presence and skin conductance as predictors. The results are reported below.

The pattern of results in the visual condition is the same as reported in the main text. With accuracy as the dependent variable, there were significant effects of pupil size, target presence, and skin conductance. Larger pupils were associated with improved accuracy (*b* = 0.155, *p* = .013), while both the presence of a target and higher skin conductance were associated with decreased accuracy (*b* = -3.676, *p* < .001; *b* = -0.119, *p* = .041).

With response as the dependent variable, there were significant effects of pupil size and target presence. Both larger pupils and the presence of a target were associated with more ‘yes’-responses (*b* = 0.135, *p* = .020; *b* = 4.188, *p* < .001). Skin conductance was not significantly related to response tendency (*b* = -0.095, *p* = .085).

In the auditory condition the pattern of results in the auditory condition is the same as reported in the main text. With accuracy as the dependent variable, there was a significant effect of target presence, such that the presence of a target was associated with decreased accuracy (*b* = -3.836, *p* < .001). The effects of pupil size and skin conductance were not significant (*b* = -0.094, *p* = .053, *b* = -0.047, *p* = .508).

With response as the dependent variable, there was a significant effect of target presence, such that the presence of a target was associated with more ‘yes’-responses (*b* = 3.884, *p* < .001). Pupil size and skin conductance were not significantly related to response tendency (*b* = -0.060, *p* = .144; *b* = -0.064, *p* = .345).

The pattern of results in the auditory condition is slightly different than in the analysis reported in the main text, whereby here the effect of pupil size on accuracy is significant, indicating that performance is worse for larger pupils.

### Correlations between explanatory variables

Figure 1 depicts the correlations between each pair of explanatory variables.

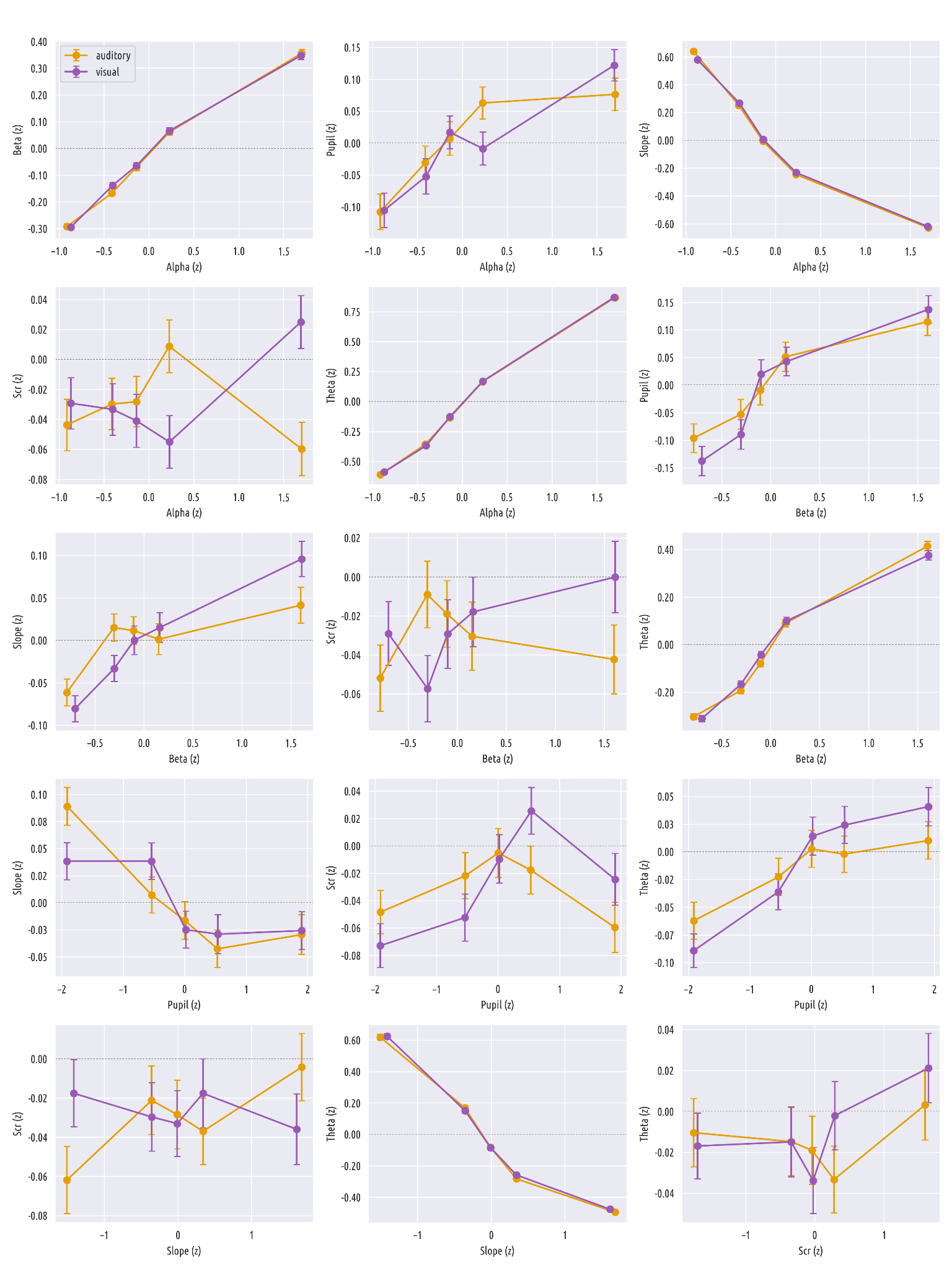

*Figure 1.* The average within-subject correlations between all pairs of explanatory variables in the visual (purple) and auditory (orange) conditions. All values are normalized. Error bars represent the standard error of the mean (SEM) within the bin.

### EEG Figures

Figures 2, 3, and 4 show the group-average power spectral density (PSD) of the analysis channels, topomaps split by frequency band, and time-frequency spectra of the prestimulus interval, for each condition separately. Note that the PSD does not include channels that were deemed bad for a subset of the subjects. The TFR was computed using Morlet wavelets across 30 linearly-spaced frequencies between 4 and 30 Hz, with the number of cycles set to half the frequency (n_cycles = f/2). Power was averaged across epochs and across channels, and cropped to the pre-stimulus window (−500 to 0 ms).

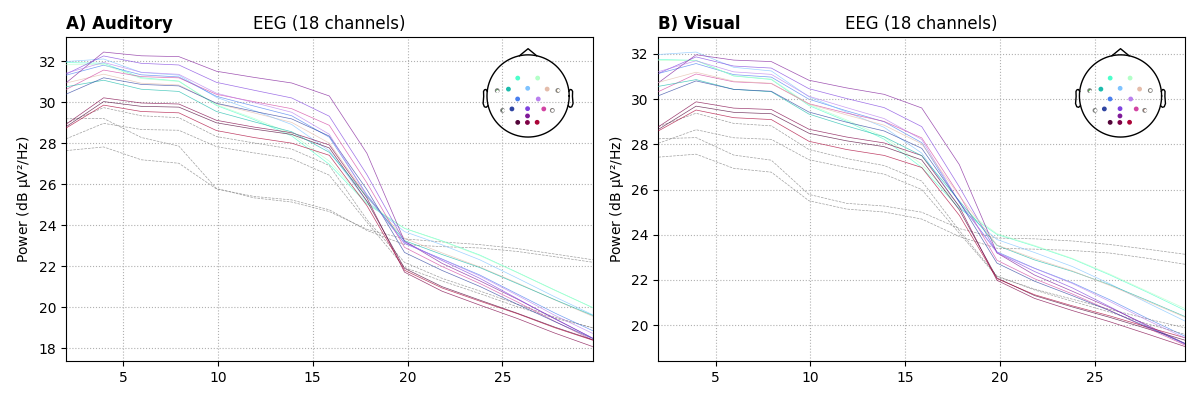

*Figure21.* PSDs showing how power in each channel is distributed across the frequency spectrum, split by condition. Using the average of all subjects’ data, including only analysis channels (O1, O2, Oz, POz, Pz, P3, P4, P7, P8, T7, T8, C3, Cz, C4) . Note that if a channel was deemed bad for any subject it was removed from the PSD (but plotted here as dashed lines). A) Auditory condition. B) Visual condition.

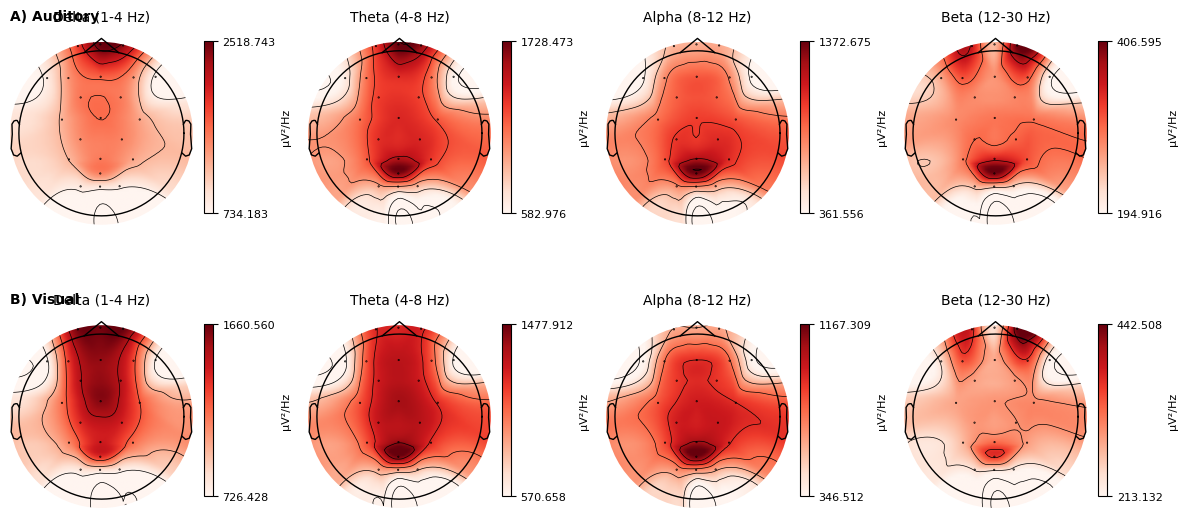

*Figure 3.* Topoplots showing which channels contribute most which frequencies. Split by condition and averaged across subjects. A) Auditory condition. B) Visual condition.

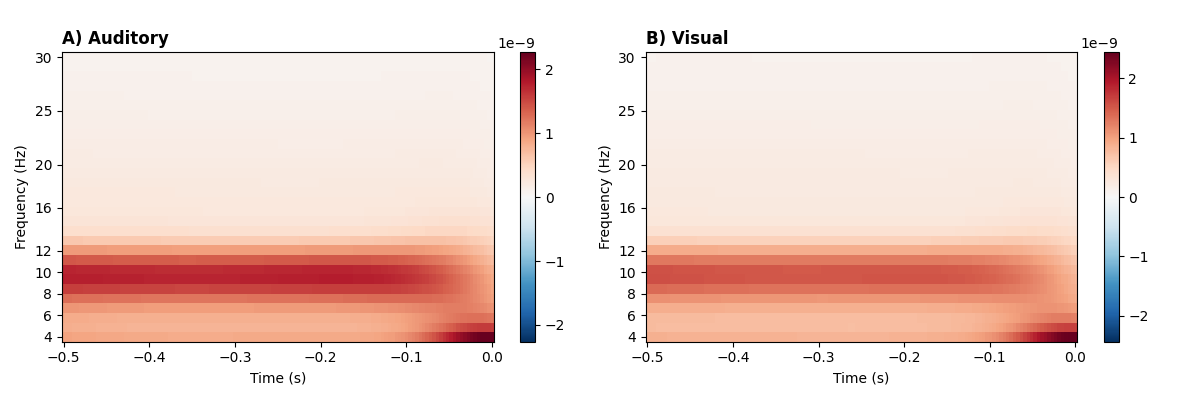

*Figure 4.* The TFR representation of the pre-stimulus interval (-500 to 0ms, split by condition. A) Auditory condition. B) Visual condition.
